# Burrow Morphology of Genus *Ocypode* (Brachyura: Decapoda: Ocypodidae) Along the Coast of Karachi

**DOI:** 10.1101/128033

**Authors:** S. Odhano, N. U. Saher

## Abstract

Burrow morphology of *Ocypode rotundata* and *O. ceratophthalma* was studied on the sandy beach of Karachi with the aim of identifying their significance and relationship to the shore environment. The small sized burrows found at low tide level and large sized burrows found at the high tide level up to dry or splash zone. The burrow count during the winter season was lower as compared summer season. Only single burrow opening was observed in *O. rotundata* and *O. ceratophthalma* oriented towards the sea. The burrow depth was between 460 to 1300 mm and 490 to 760 mm in *O. rotundata* and *O. ceratophthalma* respectively. Strong correlation (r^2^=81.2 and 89.2%) was observed between carapace length and burrow diameter of the *O. rotundata* and *O. ceratophthalma* respectively. For the grain size analysis, maximum amount of grain resulted with fine sand 57.04% (2.5Φ, 3.0Φ). For anthropogenic analysis, data showed no any significant difference (P value =0.128 and 0.671) from two sites but number of burrow counts decreases as the number of human activity increasing day by day at the selected beaches.

## INTRODUCTION

Ghost crabs of the genus *Ocypode* are so called semi-terrestrial species [6,11,19,21,23,25]. These crabs are frequently found in tropical to sub-tropical areas along the sandy coasts of the world, starting from American Atlantic through the Mediterranean, Red Sea to American Pacific and Indo-Pacific regions [11,19,27]. *O. ceratophthalma* and *O. rotundata* are commonly distributed in the Indo-Pacific region, found in large quantities above the high tide mark on sandy shores [1,6,7,10,19,20,27]. These species are prominently macroscopic invertebrates inhabiting the sandy beaches found along the coasts of Pakistan.

Ghost crabs are relatively large invertebrates typically nocturnal, but their juveniles can also be seen during day time because they do not have capability to spent much time inside their burrows. As they are nocturnal species they feed during night time. They have ability to feed on any type of food (such as, macroscopic, microscopic, live or dead animal or plant materials or sometimes they are called scavengers [26].

Those marine organisms living in soft sediments of marine coastal area have developed a burrowing adoptability which is commonly found in the invertebrates [22]. Burrows can be constructed by ghost crabs in different shapes and sizes then symbolised through their alphabetical terms such as J-, Y-, U-shaped which purely depends on sediment properties, tidal level and shore types [5, 6]. Ghost crabs can construct deep and complex burrows may be as deep as 2 m [10] which provide shelter against climatic extremes and predators and serve as sanctuary during molting and motherliness [6, 13]. The burrow openings of ghost crabs are circular with accumulated sand mounds and are often surrounded by intense feeding lines left by the crabs [5,6]. There are clearly visible entrances of a ghost crab on the surface of sandy beaches which are maintained as territory [24] by which counting of these holes can help to measure the densities of ghost crab easily [13,15].

The coastlines of the world are dominated by sandy beach ecosystems [3,14]. By increasing the population of human beings, the natural habitats of sandy beaches are being destroyed at the accelerating rate [9]. Coastal development caused by human activities effects on the extensive alterations in the coastal beaches. Ghost crabs can be used for the assessment of human impact on the beach environment [2] because they are most important part of food chain on sandy beaches.

The history shows that the species belong to the genus *Ocypode* [25] were studied comprehensively for their behaviour and physiology but studies regarding burrow morphology are limited [6, 22]. In Pakistan, the significance of ghost crab burrow morphology and relative growth analysis on existent beaches has not been explored yet. It is under the author’s knowledge that no previous work has been done so far on burrow morphology and relative growth analysis on the genus *Ocypode* [25] along the coast of Karachi, Pakistan.

## MATERIALS AND METHODS

### Study area

There is about 990 km [17,18] of the coast line of Pakistan in which the Karachi coast comprises of 100 km [18], where the detailed study was carried out. The beach of Sandspit (24° 50’N, 66° 56’E) is covered by wide areas of mangrove on its back shore. The backshore and foreshore are separated by a wide strip of road which connects the Hawksbay coastline with Manora beach [12] Sandspit in between them. The total distance from Hawksbay to Manora is about 20 km [12]. The study was carried out on two different stations of Sandspit (S1) opposite to WWF regional office and (S2) near the CEMB laboratory. These sites can easily be reachable, associated with the large population of ghost crabs. These are habitually nesting sites for green turtle *Chelonia mydas* [12].

The both stations (S1 and S2) were subdivided into the 2 localities: locality one was termed as Upper Shore Limit (USL) which was 38 feet away from the surf zone of sea and locality two was named as Lower Shore Limit (LSL) started from surf zone. The number and size distribution of burrows within each locale was examined. Total number of burrows, total number of pyramids were observed and counted before analysis.

The beach of Sandspit (24° 50’N, 66° 56’E) is covered by wide areas of mangrove on its back shore. The backshore and foreshore are separated by a wide strip of road which connects the Hawksbay coastline with Manora beach Sandspit in between them [12]. The total area of Hawksbay and Sandspit is about 20 km, where detailed study was carried out on two different stations of Sandspit [12]; area opposite to WWF regional office (S1) and area routes towards the Manora beach where the CEMB laboratory is situated (S2).

### Sampling methodology

The present study was carried out for two years from March 2011 to September 2012. Data was collected in four months (March, April, August, September) in each year (2011-2012). In each station (S1 and S2) about eight transects were placed for burrow count and cast filling through line transect method. Each transect was a 2 m^2^ in size. A measurement was taken from upper shore limit to the lower shore limit. Four quadrates were placed in upper shore limit and in lower shore limit. Aqueous solution of plaster of Paris to water with the ratio of 2:1 was poured into the selected crab burrows until the burrows were completely filled. The solution could dry for 30 to 60 minutes [16, 20]. This technique has significantly improved our awareness that how ghost crabs construct their burrows in different shapes. After pouring the cast into the burrow if the crabs emerge out were collected and placed into the marked poly bags and brought into laboratory for relative growth analysis. On several occasions the crabs trapped inside their burrows and could not emerge out because of the depth and branching of the burrow and immediate drying effect of the cast. The casts were excavated carefully as they become solidified and carefully taken to the laboratory then measurements of burrow proportions were carried. The casts were cleaned prior to measurement through brush to remove excess sediments. The complete casts were used for the further analysis. Burrows were sorted according to their concerned species and shapes. Burrow counts and cast filling were replicated during each visit, the purpose of this replication to test the human disturbance over time.

### Sediment analysis

To observe the variation in the sediment structure sediment samples were taken nearby to the casting area from the depth of 30 cm for each quadrate replicate, to determine the sediment properties. Sediment analysis was carried out in laboratory to observe the percent organic matter and grain size of sediment through the oven, furnace and sieves with different mesh sizes respectively [18]. For percent organic matter, 200-g sediment taken into pre-weighted crucibles then placed into the oven at 70°C for 5 hrs, then weighted again and positioned in a furnace at 400°C for 24 hrs after that weighted again and data was used for statistical analysis.

For the grain size analysis, sediment was dried at room temperature and treated to have a permanently wrinkled appearance. About 100 g of sediment sample were taken from a dried sample of sediment and sieved through the sieve machine by keeping standard mesh sized sieve (Fig. 5). Time was kept constant during the sieving period about 15 minutes for getting perfect results. The sediments retained on each sieve were collected and weighted then collected data was statistically analyzed to obtain the percent grain size.

### Anthropogenic impact

For the purpose to explore the anthropogenic impact on ghost crab population the involvement of human activities on sandy beaches was observed; that can greatly cause the reduction in crab density. About 20-m area from each site of Sandspit (S1 and S2) selected and one reference site selected which was most populous site and regularly visited site by humans for picnic. The ghost crab densities were calculated by counting the number of active burrow openings on the beach surface once visited the site. In each section burrows counted from the selected zone through line transect method. Measurement was taken from upper shore limit to the lower shore limit. Four lines were positioned in each limit (4 in upper shore limit and 4 in lower shore limit) and burrow densities were made as the number of burrows per line. Burrow counts were replicated during each visit, the purpose of this replication to test the human disturbance over time. This study was carried out 8 times in each year (2011 & 2012) during two times of each month (March, April, August and September).

### Statistical analysis

Following analysis was carried out by using the MS Excel (ver. 2013), Minitab ver. 17 and SPSS ver. 16: Regression, correlation with burrow structure, t-test was employed between two stations. Descriptive analysis and One-way ANOVA were employed to the data obtained during experimental work (i.e., burrow morphology, crab morphology, sediment structure and anthropogenic analysis).

The regression analysis (y = a + b x) was used for the study for morphometric analysis of each population (male and female) where (b) designates the slope and (a) as a Y – intercept. The t-test was employed to observed the difference between means of two different sites. The descriptive analysis was used for both populations’ data to investigate the basic difference and to observe the maximum minimum ratio of all variables along with their means and standard deviation. The One-way ANOVA that was employed with the supporting null hypothesis that states the populations of genus *Ocypode* from two different sites were same.

For the burrow morphology analysis following variables were used for the analysis: total burrow length (TBL), total burrow depth (TBD), opening diameter of burrow (ODB), branch (shaft) length (BL), burrow distance from sea (BDS). For relative growth analysis only 6 morphological characters were selected: carapace length (CL), carapace width (CW), enlarged chela length (EnL), enlarged chela width (EnW), abdominal length (AbL) and abdominal width (AbW). For Sediment Analysis percent of organic matter, and grain size were observed. And finally, for anthropogenic impact every visit burrows were counted from the selected area to identify the impact of humans over the burrows.

## RESULTS

### Burrow morphology

Total 107 complete burrow casts were collected in which ghost crabs *O. rotundata* 60 casts (46 males and 14 female), *O. ceratophthalma* 39 casts (28 males and 11 female) and another 8 unknown casts (Table 1, Fig. 3) were hosting. Where *O. rotundata* male occupied the highest percentage (42.99%) then the other found species and unknown as cast secured the lowest percentage (7.47%) (Table1). The species *O. rotundata* represented four different types of burrow structures: J-shaped (12 burrow casts), L-shaped (13 burrow casts), Network (3 burrow casts), and I-shaped (32 burrow casts) ((Figs. 1, 2). Whereas ghost crab *O. ceratophthalma* also characterized with four types of burrow structures but out four two were different from *O. rotundata:* C-shaped (15 burrow casts) and Y-shaped (14 burrow casts) along with two similar types of burrow casts J- and L-shaped (4 and 6 burrow casts respectively) (Figs. 1, 3). The maximum number of collected casts were straight in I-shaped (32%); while the minimum number of collected, casts were Network (3%) that were only observed in the *O. rotundata* (Fig. 1).

**Table 1.** Percentage occupied by two *Ocypode* species during burrow cast collection

| Species | Sex | Total<br>number | Percentage<br>(%) |
| --- | --- | --- | --- |
| *Ocypode rotundata* | Male | 46 | 42.99 |
|  | Female | 14 | 13.08 |
| *O. ceratophthalma* | Male | 28 | 26.16 |
|  | Female | 11 | 10.28 |
| Unknown | ??? | 08 | 7.47 |
| Total |  | 107 | 100 |

Through regression analysis burrow opening diameter and total burrow length showed positive allometric relation with carapace length in *O. rotundata* (R^2^= 81.2%) in both variables respectively. While in *O. ceratophthalma* positive allometric relation was observed in burrow opening diameter with carapace length (89.2%) while isometric relation was observed in total burrow length with carapace length (R^2^= 89.2%) (Figure 4 a-d). Other comparative variables were also used for regression analysis where carapace width was used as independent variable (Table 2).

**Table 2.** Regression analysis of comparative variables in two *Ocypode* species N= total number of individuals, R^2^=? AM= allometric growth, CL= carapace length, CW= carapace width, EnChlL= enlarged chela length, SmChlL= small chela length, AbL= abdominal length, AbW=abdominal width.

| Comparative<br>variables | Sex | N | *Ocypode rotundata* |  |  | *O. ceratophthalmus* |  |  |  |
| --- | --- | --- | --- | --- | --- | --- | --- | --- | --- |
| | | | Y=a + b(x) | $R^2$ | AM | N | Y=a + b(x) | $R^2$ | AM |
| CL vs CW | M | 14 | Y=05.02+1.00x | 0.78 | O | 28 | Y= 2.50+1.08x | 0.95 | O |
|  | F | 46 | Y=-0.41+1.16x | 0.87 |  | 11 | Y= 2.64+1.11x | 0.98 | +ve |
| CW vs | M | 14 | Y=4.03+0.72x | 0.88 | -ve | 28 | Y= 2.91+0.81x | 0.88 | O |
| EnChlL | F | 46 | Y=1.35+0.76x | 0.84 |  | 11 | Y=-4.48+0.84x | 0.88 | O |
| CW vs | M | 14 | Y=2.97+0.54x | 0.85 | -ve | 28 | Y= 4.85+0.44x | 0.69 | -ve |
| SmChlL | F | 46 | Y=3.10+0.52x | 0.84 |  | 11 | Y=-4.42+0.73x | 0.96 | O |
| CW vs AbL | M | 14 | Y=5.84 + 0.51 | 0.67 | -ve | 28 | Y= 1.02+0.54x | 0.75 | -ve |
|  | F | 46 | Y=7.19 + 0.47x | 0.68 |  | 11 | Y= 0.57+1.97x | 0.71 | +ve |
| CW vs AbW | M | 14 | Y=-2.18+0.50x | 0.50 | -ve | 28 | Y= 6.33+0.09x | 0.021 | -ve |
|  | F | 46 | Y=-0.41+0.51x | 0.59 |  | 11 | Y= 0.79+0.58x | 0.69 | -ve |

A two-sample t-test was employed on the basis of statistical hypothesis in which the mean of two populations were examined. A two-sample t-test was observed to compare whether the average difference between two populations is really significant or if it is due instead of indiscriminate chance. A significant difference (0.007 and 0.001) was observed in both species *O. ceratophthalma* and *O. rotundata* regarding carapace width (35.44±2.81 and 45.76±1.14) and carapace length (29.96±2.24 and 40.23±1.57) respectively.

The J-shaped burrows had the smallest volume with a mean opening diameter (OD) of 38.31 mm in *O. rotundata* (Fig. 2, Table 4); whereas in *O. ceratophthalma* Y-shaped burrows showed the smallest OD of 52.21 mm (Fig. 3, Table 3). The primary and secondary arms joined together into a straight shaft and ended up in a chamber at the base (Fig. 2, 3). All the primary arms faced the seaward side and vice versa for the secondary arms in *O. ceratophthalma* but not in *O. rotundata*. The highest mean opening burrow diameter in *O. rotundata* was 87.0 ± 67.7 mm along with the mean CL was 26. 00 mm; however, in *O. ceratophthalma* C-shaped burrowed 59.87±14.31 mm OD and made by crabs with mean carapace length 37.6 mm (Table 3; Fig. 2, 3).

**Table 3.**
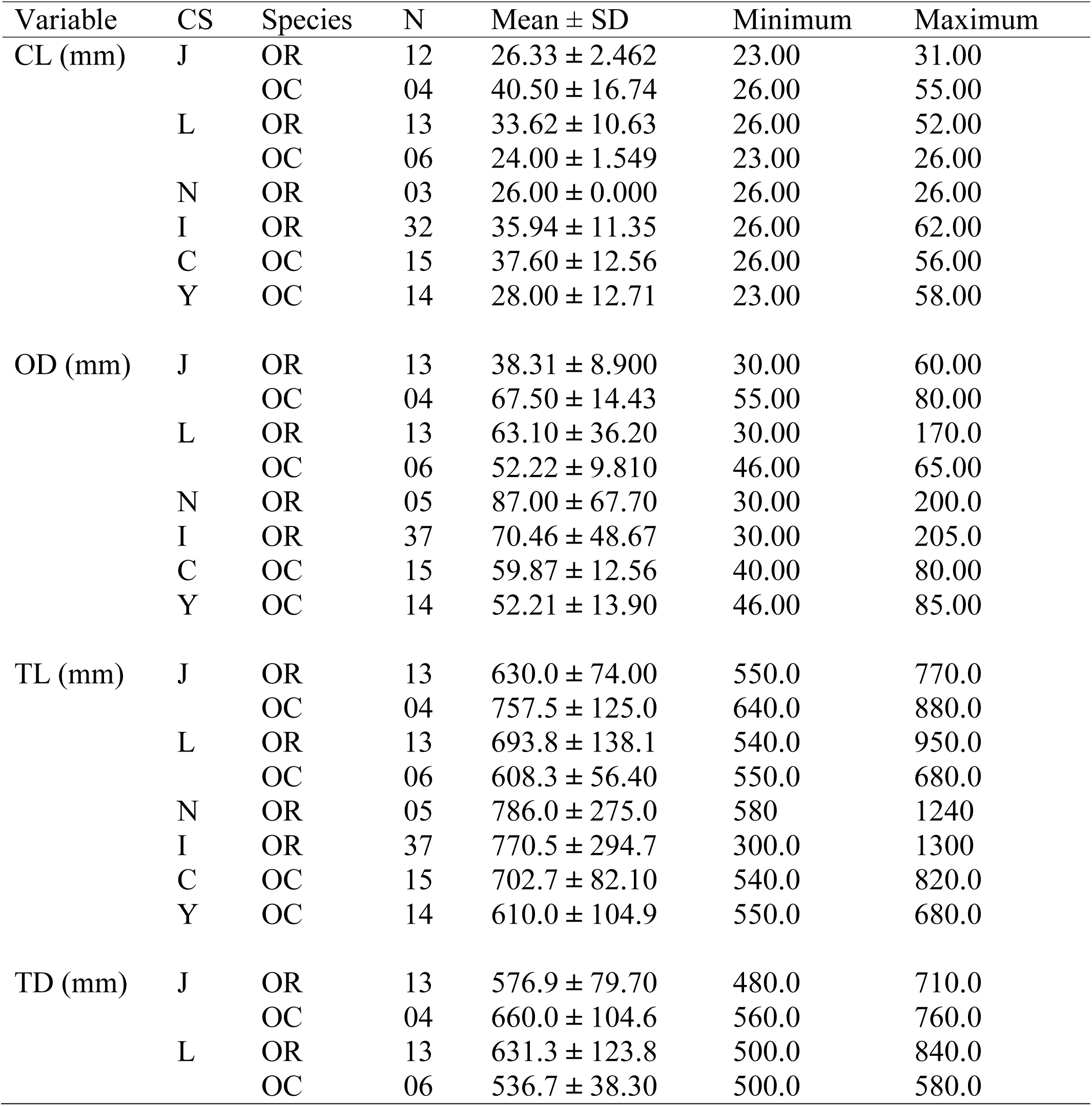
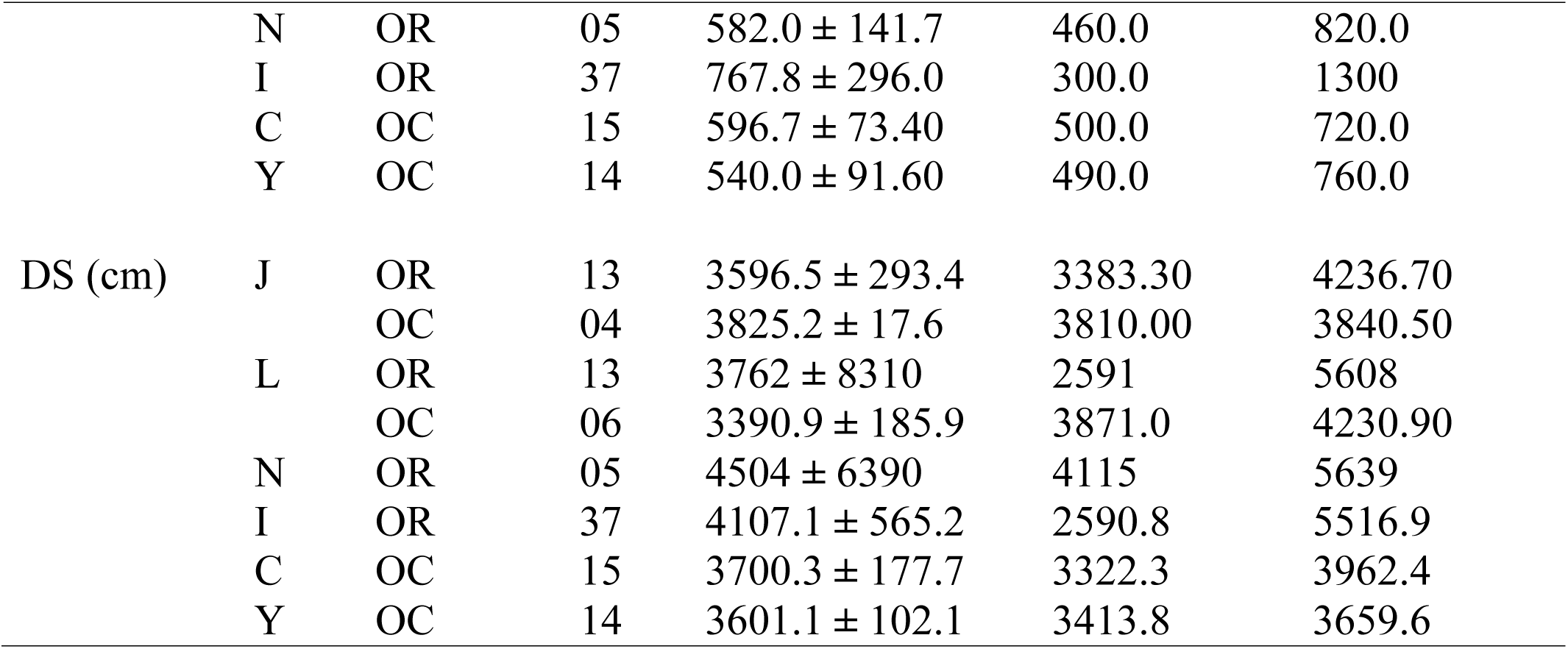
Descriptive analysis of burrow cast structure of *Ocypode* species along the coast of Karachi (OR = *O. rotundata*, OC = *O. ceratophthalma*, CL= carapace length, OD= opening burrow diameter, TL= total length of burrow, TD= total depth of burrow, CS= cast shape)

| Variable | CS | Species | N | Mean ± SD | Minimum | Maximum |
| --- | --- | --- | --- | --- | --- | --- |
| CL (mm) | J | OR | 12 | 26.33 ± 2.462 | 23.00 | 31.00 |
| | | OC | 04 | 40.50 ± 16.74 | 26.00 | 55.00 |
| | L | OR | 13 | 33.62 ± 10.63 | 26.00 | 52.00 |
| | | OC | 06 | 24.00 ± 1.549 | 23.00 | 26.00 |
| | N | OR | 03 | 26.00 ± 0.000 | 26.00 | 26.00 |
| | I | OR | 32 | 35.94 ± 11.35 | 26.00 | 62.00 |
| | C | OC | 15 | 37.60 ± 12.56 | 26.00 | 56.00 |
| | Y | OC | 14 | 28.00 ± 12.71 | 23.00 | 58.00 |
| OD (mm) | J | OR | 13 | 38.31 ± 8.900 | 30.00 | 60.00 |
| | | OC | 04 | 67.50 ± 14.43 | 55.00 | 80.00 |
| | L | OR | 13 | 63.10 ± 36.20 | 30.00 | 170.0 |
| | | OC | 06 | 52.22 ± 9.810 | 46.00 | 65.00 |
| | N | OR | 05 | 87.00 ± 67.70 | 30.00 | 200.0 |
| | I | OR | 37 | 70.46 ± 48.67 | 30.00 | 205.0 |
| | C | OC | 15 | 59.87 ± 12.56 | 40.00 | 80.00 |
| | Y | OC | 14 | 52.21 ± 13.90 | 46.00 | 85.00 |
| TL (mm) | J | OR | 13 | 630.0 ± 74.00 | 550.0 | 770.0 |
| | | OC | 04 | 757.5 ± 125.0 | 640.0 | 880.0 |
| | L | OR | 13 | 693.8 ± 138.1 | 540.0 | 950.0 |
| | | OC | 06 | 608.3 ± 56.40 | 550.0 | 680.0 |
| | N | OR | 05 | 786.0 ± 275.0 | 580 | 1240 |
| | I | OR | 37 | 770.5 ± 294.7 | 300.0 | 1300 |
| | C | OC | 15 | 702.7 ± 82.10 | 540.0 | 820.0 |
| | Y | OC | 14 | 610.0 ± 104.9 | 550.0 | 680.0 |
| TD (mm) | J | OR | 13 | 576.9 ± 79.70 | 480.0 | 710.0 |
| | | OC | 04 | 660.0 ± 104.6 | 560.0 | 760.0 |
| | L | OR | 13 | 631.3 ± 123.8 | 500.0 | 840.0 |
| | | OC | 06 | 536.7 ± 38.30 | 500.0 | 580.0 |
|  | N | OR | 05 | 582.0 ± 141.7 | 460.0 | 820.0 |
|  | I | OR | 37 | 767.8 ± 296.0 | 300.0 | 1300 |
|  | C | OC | 15 | 596.7 ± 73.40 | 500.0 | 720.0 |
|  | Y | OC | 14 | 540.0 ± 91.60 | 490.0 | 760.0 |
| DS (cm) | J | OR | 13 | 3596.5 ± 293.4 | 3383.30 | 4236.70 |
|  |  | OC | 04 | 3825.2 ± 17.6 | 3810.00 | 3840.50 |
|  | L | OR | 13 | 3762 ± 8310 | 2591 | 5608 |
|  |  | OC | 06 | 3390.9 ± 185.9 | 3871.0 | 4230.90 |
|  | N | OR | 05 | 4504 ± 6390 | 4115 | 5639 |
|  | I | OR | 37 | 4107.1 ± 565.2 | 2590.8 | 5516.9 |
|  | C | OC | 15 | 3700.3 ± 177.7 | 3322.3 | 3962.4 |
|  | Y | OC | 14 | 3601.1 ± 102.1 | 3413.8 | 3659.6 |

**Table 4.** Detailed descriptive analysis of t annual percent of organic matter and grain size analysis

| Core | Year | Station | Organic matter (%) | Coarse slit (>4.0/230) | Very fine sand (=4.0/230) | Fine sand (=3.0/120) | Fine sand (=2.5/80) | Medium sand (=2.0/65) | Medium sand (=1.5/45) |
| --- | --- | --- | --- | --- | --- | --- | --- | --- | --- |
| Core 1 | 2011 | Shore Lab | 8.68 | 1.25 | 3.52 | 43.62 | 22.11 | 15.73 | 14.00 |
|  | 2012 |  | 7.23 | 1.15 | 2.98 | 44.62 | 21.30 | 14.63 | 15.32 |
| Core 2 | 2011 | Shore Lab | 3.93 | 1.36 | 4.37 | 43.02 | 25.07 | 13.67 | 12.03 |
|  | 2012 |  | 9.55 | 1.46 | 4.88 | 41.44 | 29.56 | 11.33 | 11.33 |
| Core 3 | 2011 | Shore Lab | 3.74 | 0.34 | 1.28 | 18.92 | 24.34 | 15.35 | 37.45 |
|  | 2012 |  | 1.86 | 1.14 | 2.18 | 38.22 | 18.66 | 17.66 | 22.14 |
| Core 4 | 2011 | Shore Lab | 5.21 | 1.07 | 3.99 | 35.84 | 33.85 | 14.75 | 12.7 |
|  | 2012 |  | 3.30 | 1.34 | 4.80 | 28.99 | 32.55 | 19.88 | 12.44 |
| Core 5 | 2011 | Shore Lab | 1.91 | 1.29 | 4.01 | 31.23 | 33.03 | 16.01 | 15.35 |
|  | 2012 |  | 3.28 | 0.66 | 3.55 | 38.55 | 24.78 | 14.99 | 17.47 |
| Core 6 | 2011 | Shore Lab | 1.26 | 1.90 | 5.03 | 40.27 | 36.58 | 11.43 | 7.12 |
|  | 2012 |  | 5.49 | 1.99 | 4.66 | 41.29 | 33.76 | 10.44 | 7.86 |
| Core 7 | 2011 | WWF Site | 1.23 | 1.75 | 4.52 | 42.62 | 21.11 | 14.73 | 15.21 |
|  | 2012 |  | 0.21 | 1.20 | 4.38 | 43.22 | 20.20 | 15.10 | 15.90 |
| Core 8 | 2011 | WWF Site | 4.47 | 1.56 | 3.37 | 41.02 | 22.07 | 16.67 | 15.03 |
|  | 2012 |  | 1.86 | 1.26 | 3.25 | 40.44 | 23.66 | 15.10 | 16.29 |
| Core 9 | 2011 | WWF Site | 1.86 | 1.34 | 2.28 | 35.92 | 29.34 | 15.35 | 16.45 |
|  | 2012 |  | 3.30 | 1.15 | 1.98 | 34.29 | 28.66 | 16.63 | 17.29 |
| Core 10 | 2011 | WWF Site | 3.30 | 2.07 | 2.99 | 31.84 | 30.85 | 16.75 | 15.7 |
|  | 2012 |  | 6.50 | 1.98 | 3.50 | 30.22 | 29.88 | 17.88 | 16.54 |
| Core 11 | 2011 | WWF Site | 1.65 | 1.90 | 4.01 | 36.23 | 28.03 | 16.01 | 13.35 |
|  | 2012 |  | 1.65 | 2.14 | 3.70 | 38.44 | 25.77 | 14.66 | 15.29 |
| Core 12 | 2011 | WWF Site | 0.21 | 0.9 | 3.03 | 40.27 | 36.58 | 15.43 | 4.12 |
|  | 2012 |  | 1.23 | 1.25 | 2.78 | 39.55 | 33.27 | 14.28 | 8.87 |

The maximum (62 mm) of carapace length was observed from the I-shaped cast; while Network shaped cast was observed (26.00 mm) (Table 3, Fig. 1). *O. ceratophthalma* showed the maximum number of the cast structure in C- and Y- shaped with 15 and 14 casts respectively (Fig. 1, Table 3). Both cast structure (C and Y) showed a maximum number of carapace length with (56 and 58 mm) (Table 3). The male species of *O. rotundata* was lacking C and Y shaped burrow structures while *O. ceratophthalma* male was missing the I-shaped and Network type of burrows. While female of *O. rotundata* species showed only I- and J-shaped structures only C- and J-shaped structures were observed in female of *O. ceratophthalma* (Table 3).

The burrow opening diameter (OD) showed the positive correlation with carapace length (CL) (R^2^= 81.2, 89.2%) of crabs *O. rotundata* and *O. ceratophthalma* respectively while total length of burrow (TL) showed positive correlation with CL (R^2^= 81.2%), in *O. rotundata* as compared to *O. ceratophthalma* where total length of burrow (TL) showed isometric correlation with carapace length (R^2^= 72.5%) (Fig. 4 a-d). A comparison of the morphological parameters of the sampled burrows reveals the contrasting burrow architecture between two species of *Ocypode*. These findings indicate that morphological variations in these burrows may enable the differentiation of species habitat in the site.

### Sediment analysis

Sediment analysis was carried out to obtain the percent organic matter and percent grain size. The yearly grain size analysis results the maximum amount of fine sand observed from both sites among which fine sand observed to be about 70% of total sand from core 6 (C6). The minimum amount coarse slit sand not more than 2% from any core, site or year was observedб details given in the figure 4 and 5. This analysis showed the significant difference during the period (Fig. 5, Table 4). The grain size analysis showed that the maximum amount of grain found in fine sand of two sieve 57.04% (2.5Φ, 3.0Φ).

### Anthropogenic impact

Both selected sites were observed to be lower impact of human population for bathing and picnicking due to construction of private huts throughout the coastal belt. But the number of people increased during the weekend days (Saturday and Sunday). Both sites are morphologically similar, burrow density was analysed through One-way ANOVA and descriptive analysis was used to express the difference between the two sites. The analysis showed that the distribution of burrow densities varied considerably with grades of human disturbance (S1 and S2) but no any significant difference was observed in positions across the shore as shown in Table 5. The reason for not showing any significant difference that the selected sites were not frequently visited by public due to private huts which are constructed throughout the coastline which cause the public restricted area for bathing and picnicking which showed a significant difference (P =0.008) which was used as reference site. Importantly distribution of burrow densities showed variation with sequential changes during every visit which describes that number of burrow densities change in space and time which may lead to the cause of anthropogenic impact. The data showed (Table 5) number of burrows decrease by the passage of time due to increased number of human activity over the beach.

**Table 5.** Decriptive analysis of total number of burrows showing anthropogenic impact. Minimum number of burrows, maximum number of burrows and One-way ANOVA P value of two selected sites along with Picnic site ()

| Site | Year | Total | Mean±SD | Minimum | Maximum | P |
| --- | --- | --- | --- | --- | --- | --- |
| Picnic site | 2011 | 103.00 | 12.88±4.70 | 8.00 | 21.00 | 0.008 |
|  | 2012 | 60.000 | 7.500±1.773 | 5.000 | 10.000 |  |
| Shore Labe | 2011 | 1320.0 | 165.0±47.0 | 69.0 | 188.0 | 0.128 |
|  | 2012 | 987.0 | 123.4±38.5 | 78.0 | 226.0 |  |
| WWf Site | 2011 | 1161.0 | 145.1±32.1 | 103.0 | 192.0 | 0.671 |
|  | 2012 | 1400.0 | 175.0±41.2 | 83.0 | 210.0 |  |

## DISCUSSION

The identification of these two species by the current study was featured by using the available identification keys and these both species were commonly found abundantly along the both selected stations. Both stations were showing major difference in terms of their tidal height and sediment properties (Fig. 2) and most importantly, human impact that is also the major factor on and construction of their burrows. Both the areas are separated by picnic point (Sandspit picnic point) where a little or no any ghost crab population was observed. The ghost crab species *O. rotundata* and *O. ceratophthalma* showing sympatric relationship is widely distributed along the sandy beaches of Pakistan on high tide mark.

Total 106 casts were observed, which shows that most casts were collected *O. rotundata* 60 species (55.66%) while the *O. ceratophthalma* 39 species (36.79%) remaining 8 species (7.55%) were unknown casts where no occupants were observed (Table 3).

When the cast structure was compared with these two species I-shaped and N-shaped structures were only found in *O. rotundata* and C- and Y-shaped structure were found in *O. ceratophthalma* which clearly distinguishes the two-different species according to their burrow structure. Now these species can easily be identified according to their burrow structure.

Space and food are two fundamental resources required by organisms that provided by the sediment which supports the predominantly space for the burrowing and deposited food for organisms. Particle size can also regulate the dispersal and stratification of crabs by manipulating the organic matter that the grainier sediments usually have greater amount of organic content then other type of sediment [17]. Sediment percent grain size showed 70 to 95% sediment were fine to medium sand at both stations. The dispersal of particle sizes within a substratum is designated by sorting and skewness features [4]. The sorting co-efficient can be classified as moderately well sorted to moderately sorted at both stations.

Sandspit beach comprises of ghost crabs are dominant and can be placed as top carnivores. This is the first report of burrow morphology and distribution of ghost crabs along the coastal areas of Pakistan. Ghost crabs inhabit the vast intertidal zone of almost all sandy areas of world frequently tropical to sub-tropical areas.

The ghost crabs are observed to be found on fine to medium grain sized sediment. The ghost crab density, distribution and zonation can be influence by many other factors such as competition (interspecific or intraspecific) along with temperature and light [8].

